# Enriched sleep environments lengthen lemur sleep duration

**DOI:** 10.1101/2021.06.02.446761

**Authors:** Alexander Q. Vining, Charles L. Nunn, David R. Samson

## Abstract

Characteristics of the sleep-site are thought to influence the quality and duration of primate sleep, yet only a handful of studies have investigated these links experimentally. Using actigraphy and infrared videography, we quantified sleep in four lemur species (*Eulemur coronatus, Lemur catta, Propithecus coquereli*, and *Varecia rubra*) under two different experimental conditions at the Duke Lemur Center (DLC) in Durham, NC, USA. Individuals from each species underwent three weeks of simultaneous testing to investigate the hypothesis that comfort level of the sleep-site influences sleep. We obtained baseline data on normal sleep, and then, in a pair-wise study design, we compared the daily sleep times of individuals in simultaneous experiments of sleep-site enrichment and sleep-site impoverishment. Over 163 24-hour periods from 8 individuals (2 of each species), we found strong evidence that enriched sleep-sites increased daily sleep times of lemurs, with an average increase of thirty-one minutes. The effect of sleep-site impoverishment was small and not statistically significant. We conclude that properties of a sleep-site enhancing softness or insulation, more than the factors of surface area or stability, influence lemur sleep, with implications regarding the importance of nest building in primate evolution and the welfare and management of captive lemurs.

## Introduction

Sleep is a period of behavioral quiescence and reduced responsiveness to external stimuli, thus making it a vulnerable and dangerous state [1]. Consistent with this observation, comparative studies have revealed that risk of predation at the sleep-site covaries negatively with sleep duration across mammals [2–4]. Sleep-sites likely also vary in quality and security along dimensions that involve the physical comfort of the substrate, level of concealment from conspecifics and predators, and thermal properties [5–7]. A major question concerns how characteristics of the sleep site influence sleep quality, and how that in turn influences organismal function and fitness.

Considerable effort has been put into studying the role of sleep-site characteristics for apes. The *sleep quality hypothesis*, for example, proposes that apes construct beds to permit less fragmented, undisturbed sleep that promotes sleep quality through either greater sleep intensity [deeper slow-wave sleep (SWS) and/or REM sleep] or longer individual sleep stages [8–11]. This hypothesis has been supported by increased effort chimpazees put into building more complex nests [12] and observations of orangutans (*Pongo pygmaeus wurmbii*) in Southern Borneo selecting sleep sites for comfort and stability rather than predator defence [13]. More recently, comparative analysis has shown that captive orangutans (*Pongo spp*.) are characterized by deeper, more efficient sleep than baboons (*Papio papio*.) [14]. Moreover, experimental work has demonstrated that orangutans exhibit higher quality sleep with less gross-motor movements and greater overall sleep times when using complex sleeping platforms [15]. For wild, individually sleeping apes, sleep-site modifications may improve sleep through several mechanisms. The removal or covering of protruding substrate reduces stress on tissues [12]. Enlarged surface area and functional concavity of nests also obviate the need to adjust posture during sleep to prevent falls.

Other primates exhibit a wide range of sleep-site selection behaviors, but do not build daily nests as do apes [16]. Lemurs, when they do nest, appear to do so more for predator defense and infant care than for comfort, and they do not conduct nightly sleep site modifications as great apes do – making their nests functionally more akin to those of birds than apes [6]. Though these cladistic differences in sleep-site modification behavior are likely explained in part by advanced cognition and social learning in apes [5,17], sleep-site comfort may also have less functional impact on lemurs and monkeys.

Here, we investigate the effect of sleep-site characteristics on sleep-wake activity and duration in four species of lemur (*Eulemur coronatus, Lemur catta, Propithecus coquereli*, and *Varecia rubra*) by experimentally enriching and impoverishing sleep-sites in pairs of captive lemurs simultaneously. We experimentally tested the hypothesis that the comfort and stability of the sleep-site influences lemur sleep duration. Based on this hypothesis, we predicted that 1) enriching a sleep-site with soft, insulating materials would increase sleep duration and 2) impoverishing a sleep-site by removing flat, stable, above-ground surfaces would decrease total sleep time. In previous work, we found that disrupted sleep influences aspects of lemur behavior and cognition [18]. In wild grey mouse lemurs, similar cognitive tests predicted body condition and survival [19], suggesting that sleep-related changes have functional consequences. Thus, in addition to implications for animal welfare, understanding the links between sleep sites and sleep may also inform understanding of primate behavior, ecology and evolution.

## Methods

### Study subjects

Research was performed at the Duke Lemur Center (DLC), in Durham, North Carolina, USA, where subjects were housed in dyadic, sex-balanced groups (i.e., with one male and one female). We generated actigraphic data from eight individuals, with one male and one female from each species. For detailed biographical information on the study subjects, see Bray et al. [20]. Animals received unlimited access to water and fresh fruit, vegetables, and Purina monkey chow on a daily basis. Animal use and methods were approved by the Duke University Institutional Animal Care and Use committee (Protocol #: A236-13-09) and the DLC Research Committee.

### Data collection

The study was conducted over four months from May to August in 2015. Given each experimental protocol involved the work of multiple DLC staff and researchers, the research needed to be distributed on a per species basis over multiple months. This effort generated a dataset of 40-42 nights per species, totaling 163 twenty-four-hour periods, with circadian activity continuously recorded using MotionWatch 8 (CamNtech) tri-axial accelerometers. These actigraphic sensors are lightweight (7 g) and attached to standard nylon pet collars. Animals were monitored to ensure no adverse reactions to the collar; subjects acclimated to the collars within 2 hours and wore them throughout the study. The study took place in indoor housing only to control for temperature and light conditions.

Using actigraphy and videography, response variables were generated from processed activity logs recorded at one-minute epochs. Recent advances in scoring algorithms have increased accuracy in detecting wake-sleep states and total sleep times [21]. Using actigraphy data, we generated *twenty-four-hour total sleep times* (TTST) for individuals in each species. We followed protocols used in previous primate sleep studies [22–26] and in prior work by our group performed at the DLC [20,27]. The sensor sampled movement once per second at 50 Hz and assigned an activity value per one-minute epoch. We used the operational definition of behavioral sleep measured via actigraphy as the absence of any force in any direction during the measuring period [28]. Following procedures in our previous research [20], we determined that animals were consistently at rest, and thus inferred sleep states (i.e., sustained quiescence in a species-specific posture), when actigraphy values were less than or equal to four. To validate sleep states [29], we also obtained infrared videography data (AXIS P3364-LVE Network Camera).

### Experimental procedure

In a pair-wise experimental design, individuals from each species underwent two weeks of simultaneous testing after being measured in a baseline condition. During week one (baseline), each pair experienced normal sleep conditions. These conditions were the same as normal indoor operating protocol for the DLC, where individuals had access to two housing cells with mounted, raised shelves and basic amenities (e.g., raised square crates attached to the walls). To reduce the confounds of outdoor lighting and temperature, the baseline conditions only differed from normal conditions in that individuals were restricted to sleeping indoors and did not have access to the outside enclosure during the nighttime period. Baseline differed from the experimental conditions in the options for sleep, with animals allowed to sleep socially (in pairs) during baseline but separated at night in the experiments.

During week two – the beginning of the pair-wise experiment – an individual was randomly chosen for one of two experimental conditions: *sleep enrichment* or *sleep impoverishment*. The other individual of the pair underwent the opposite treatment. One treatment was applied to each of the two housing cells the lemur pair had access to during baseline, and subjects restricted to their respective housing cell following evening caretaking. The third week was a reversal of the previous week’s condition, with individuals switching sleep-sites and thus experiencing the opposite condition. During experimental nights, individuals were isolated from the pair-mate, but they were in vocal communication and could touch one another between the enclosures, thus providing some of the typical social conditions they experienced in baseline conditions. This was done to reduce the influence of separation on individual level sleep-wake expression.

Housing cells were enriched by providing lemurs with a high-quality plastic sleeping box (30 cm × 60 cm) with open slats on the side to permit ventilation; additionally, a 2.5 cm slab memory foam mattress was embedded in the base of the box with a small nylon blanket placed on the top of the mattress (Figure 1). Housing cells were impoverished by removing all enrichments items (including sleeping crates available in baseline conditions) and removing normally available wall shelves, leaving only narrow ledges for above ground perching during sleep.

**Figure 1.**
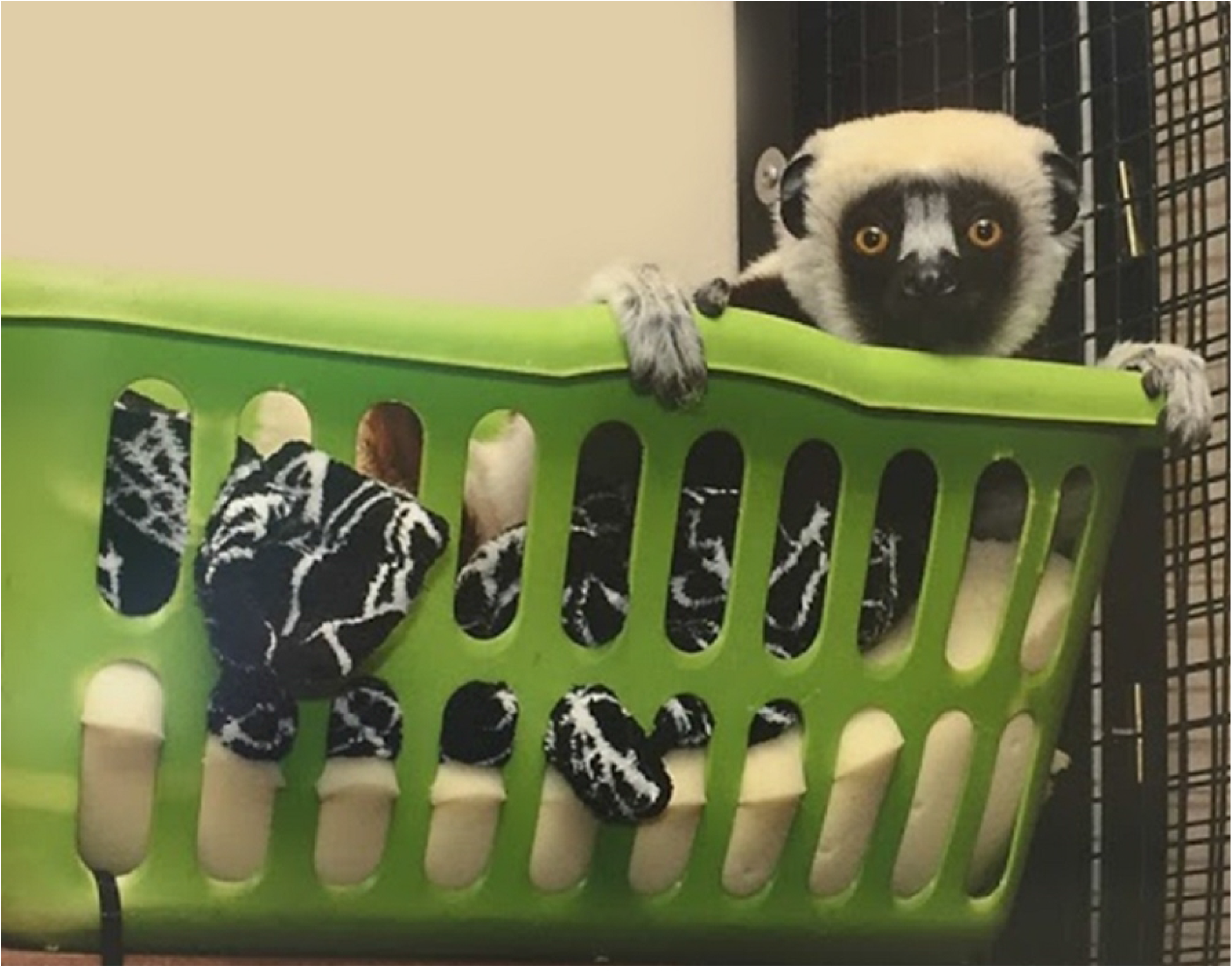
The sleep enrichment experimental condition. A *Propithecus coquereli* subject perched in the enriched sleep-site provided in the sleep enrichment experimental condition.

We used infrared videography to determine whether the subject in the sleep enrichment condition was using the sleeping box. We removed one 24-hour period from the analysis for an individual who failed to use the box on one night. In all conditions, the nighttime period was considered to start at 18:00 and end at 06:00. During the day, enrichment items were returned to the animals and they were allowed to move freely between cells.

### Data analysis

We generated descriptive statistics characterizing the distribution of inferred sleep time throughout the 24-hour period by individual and across the different experimental conditions. To test whether our experimental manipulations of sleep-site condition had a meaningful effect on lemur sleep times, we followed the approach of Pinhiero & Bates [30], building and testing a series of nested linear mixed-effects models. We began by modeling only the random effect of individual (nested within species), thus establishing a reference model that acknowledges sleep times are individually variable but assumes our experimental manipulations had no effect (Model 0). Because sleep is a biorhythm and likely to contain temporal dependencies, we expanded the reference model to include within-subject temporal autocorrelation in sleep time across nights. Based on the autocorrelation function of our sleep data, we compared Model 0 to a first order autoregressive model of TTST within subjects (Model 1). Finally, we tested the null hypothesis that our experimental manipulations had no effect on TTST by adding to Model 1 parameters describing our experimental structure. These parameters were the coefficients for the three levels of experimental condition, the two orders the conditions were presented in, and their interactions (Model 2), resulting in the equation

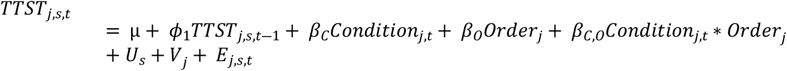

where *TTST_j,s,t_* is the TTST of individual *j* of species *s* on day *t*, *μ* is the intercept (the baseline TTST for subjects presented with the enrichment condition prior to the impoverishment condition), *ϕ*_1_ is the magnitude of the first order temporal auto-regression, *β_C_* is the regression coefficient for the experimental condition given by *Condition_j,t_*, *β_O_* is the regression coefficient for the order of experimental conditions (enrichment first versus impoverishment first) given by *Order_j_*, *β_C,O_* is the regression coefficient for the interaction of the condition-order pair given by *Condition_j,t_*\**Order_j_*, *U_s_* is a random intercept for species *s*, *V_j_* is a random intercept for individual *j*, *E_s,j,t_* is an error term, and the latter three terms are assumed to follow normal distributions with mean 0 and variances 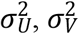, and 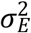, respectively.

We used delta AICs at each step to assess whether the more complex model provided a sufficiently improved fit to the data. Before making inferences about the effects of our experimental structure, we plotted the normalized residuals of Model 2 against 1) predicted values and 2) the quantiles of a standard normal distribution to conduct diagnostic checks of our model assumptions. Concluding that Model 2 met all necessary assumptions, we calculated the contrasts of each level of experimental condition (including baseline and marginal to order) and tested for significant differences in TTST between the baseline condition and the two experimental conditions using ANOVA. Finally, we calculated the intra-class correlation coefficients (ICCs) to compare the proportion of unstructured variance in our data attributable to species and individuals. We fit all models to our data using the function lme of the library nlme v3.1 (Pinheiro et al., 2007) in R version 4.0.4 (R Core Team, 2020). Our analysis can be fully reproduced using code and documentation reported in the supplementary material (S1 Analysis).

## Results

Individual sleep times are presented in Figure 2 and summarized by condition in Table 1. AIC revealed that Model 2 is the preferred model (Model 2 AIC = 1742; Model 1 ΔAIC = 4.68; Model 0 ΔAIC = 8.56). The first order autoregressive effect in this model is small (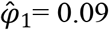, 95% CI [−0.08, 0.26]). The estimated contrast of TTST between the enriched condition and the baseline condition (marginal to order) is 31.0 minutes with a standard error of 9.03 and the hypothesis of no difference between enriched and baseline can be ruled out (*Enriched - Baseline*: df = 150, F-value = 11.4, p < 0.001). The contrast between the impoverished and baseline condition is not significantly different from 0 (*Impoverished - Baseline*: mean = 0.666 minutes, SE = 9.07, df = 150, F-value = 0.005 p = 0.943). The majority of unstructured variance in TTST given Model 2 is attributable to individual differences (ICC_subject_ = 0.675), most of the remaining variance to the error term (ICC_Error_ = 0.290), and only a small proportion to species level differences (ICC_Species_ = 0.034).

**Figure 2.**
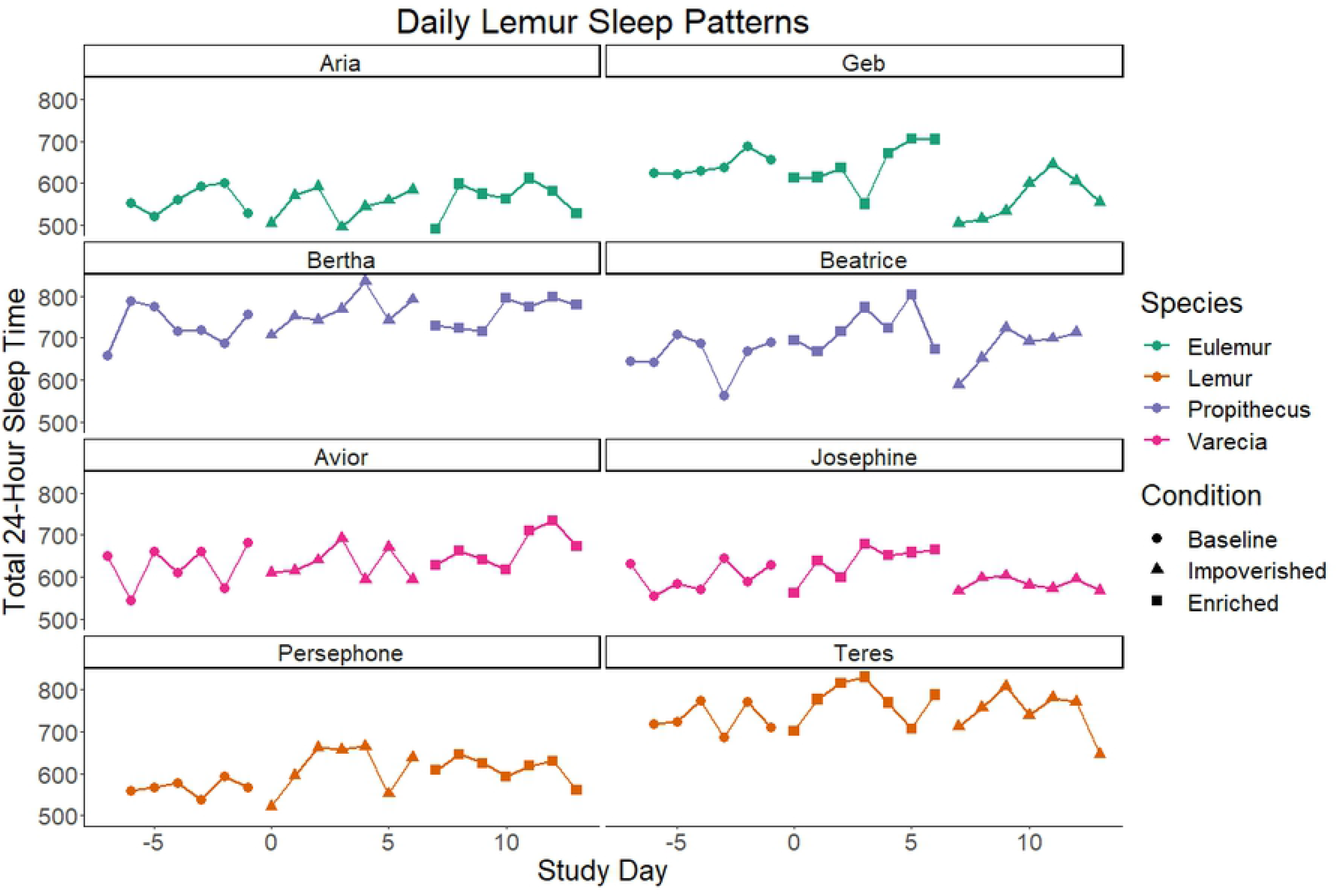
Daily lemur sleep patterns. Total sleep time in minutes for each observed 24-hour period, plotted by individual across days. Day 0 represents the start of the first experimental condition, negative values represent baseline observations. Lemurs exposed to the impoverished condition first are presented on the left, those exposed to the enriched condition first on the right.

**Table 1.** Summarized sleep times by individual and experimental condition. Mean sleep times per 24-hour period (minutes), and standard deviations (minutes) for each study individual. Base = Baseline, Enr = Enriched, Imp = Impoverished. All values have a sample size of 7, except Aria, Geb, Persophone, and Teres in the Baseline condition and Beatrice in the Enriched condition, which have a sample size of 6.

| ID | Sex | Species | Base_Mean | Base_sd | Enr_Mean | Enr_sd | Imp_Mean | Imp_sd |
| --- | --- | --- | --- | --- | --- | --- | --- | --- |
| Aria | F | *E. coronatus* | 560 | 32 | 563 | 42 | 550 | 38 |
| Geb | M | *E. coronatus* | 643 | 26 | 642 | 56 | 565 | 53 |
| Bertha | F | *P. coquereli* | 729 | 47 | 759 | 35 | 763 | 41 |
| Beatrice | M | *P. coquereli* | 658 | 48 | 721 | 51 | 678 | 50 |
| Avior | M | *V. rubra* | 626 | 51 | 666 | 43 | 631 | 39 |
| Josephine | F | *V. rubra* | 601 | 34 | 636 | 41 | 583 | 15 |
| Persephone | F | *L. catta* | 568 | 19 | 611 | 28 | 613 | 58 |
| Teres | M | *L. catta* | 731 | 35 | 769 | 49 | 745 | 54 |

## Discussion

The results of our study provide strong evidence that, for a limited subset of lemur species that do not nest outside the context of infant care, provisioning sleep sites with soft and insulated materials lengthens daily sleep durations. Among the models which we compared, the one that explicitly accounted for experimental manipulations was the most likely given our data. More importantly, we can conclude with confidence from this model that sleep site enrichment adds, on average, between 22 and 40 minutes of daily sleep relative to the baseline conditions (accounting for any effects of the order experimental conditions were presented in).

Interestingly, the manipulations of our impoverishment condition had no discernable impact on daily sleep times of our lemur subjects. Individuals in this condition were typically observed sleeping on narrow window ledges in their enclosures rather than on the ground. Though we cannot say for certain that the surface area – and hence stability – of lemur sleep sites do not affect sleep (we simply fail to reject this hypothesis), the differing impact of our impoverishment and enrichment conditions allows for some important inferences. First, we infer that the presence of soft, insulating materials is *more* important to our subjects’ sleep duration than the size of the sleeping substrate. Second, it allows us to treat the impoverishment condition as a control, indicating that incidental effects common to both manipulations, such as the separation of lemur pairs at night or the disruption of typical caretaking routines, are not sufficient to explain the differences between the enriched and baseline conditions.

Given that lemurs in our study experienced overall greater sleep duration in the enriched sleeping condition, it may have been that the sleep architecture of ancestral fixed-point sleeping primates was deep and high quality, whereas later large bodied primates that were branch sleepers traded deep, safe sleep for other advantages. There are many potential trade-offs that could result in the use of less comfortable sleep sites, despite the evidence we found that such sites could reduce total sleep times. These include predator defense, thermoregulation, parasite avoidance, group cohesion, and resource distribution [5,33]. The role of predation may be particularly important for lemurs, all of which are highly predated by raptors, boas, and fossa [34]. The high number of cathemeral (active throughout the 24-hour day) lemur species is hypothesized to result from the similarly 24-hour threat of these predator species [35]. Similarly, high predation threat may help explain why lemurs, but not other Malagasy mammals, show marked increases in female to male body size ratios relative to their mainland sister clades [36]. Thus, the anti-predator benefits that large body size and/or social living provide throughout daily activity cycles may have outweighed the benefits of secure, comfortable sleeping sites such as tree-cavities.

While large body size and social living may relax the need for secure sleep sites, additional factors are necessary to explain the benefits of flexible sleep-site selection over fixed-point nesting. Proximity to food resources has been shown to influence primate nest-construction and sleep-sites [33,37,38]. In landscapes with dynamic resource distributions, flexible sleep-sites could optimize foraging efficiency by reducing daily commutes to resource hotspots that last more than a day. Similarly, sleep-site flexibility may allow adaptive responses to changes in predation risk; why invest in sleep-site modification if you are likely to be chased off by a predator at any time? Sleeping areas can also become contaminated with parasites, suggesting that exposure to parasites might be a cost of reusing sleep-sites [39]. Conversely, certain sleeping sites can provide parasite-related benefits to primates. In the context of vector-borne diseases, for example, an enclosed site may help to obscure cues that mosquitoes use to locate hosts. Supporting this hypothesis, malaria prevalence decreases in species of New World monkeys that sleep in enclosed microhabitats [40]. Factors such as group size and body mass likely limit the ability of some primate species to use such habitats and, in general, to obtain consolidated sleep [41].

The preference our subjects showed for sleeping in enriched sites (relative to the rest of their enclosure), and the positive impact of these sites on sleep duration suggests comfort is important for sleep quality in lemurs. Likewise, the choice of our subjects to sleep on narrow, elevated ledges rather than the ground during the impoverishment condition suggests an important role of height and security in sleep-site selection, though we cannot assess the impact this has on sleep time. The dearth of nesting behavior in our subject species’ wild conspecifics further indicates that these species have evolved larger body size and/or sleep-site flexibility at the expense of sleep quality - though further data on the sleep quality of smaller lemurs sleeping in fixed-point nests or tree cavities will be essential for fully testing this hypothesis. Thus, sleeping sites have likely been a driver of sleep duration and sleep-wake regulation throughout primate evolution, but the benefits of sleep quality have not always been the predominant factor in sleep site selection.

Only with the emergence of frequent, secure platform construction in ancestral apes has deep, REM heavy sleep architecture (re)-emerged, expressed in its most extreme case in humans [42]. More recently, sleeping on the ground in sentinalized groups may have permitted even higher quality sleep along the human lineage [43]. Cognitive ability likely explains some differences in nesting behavior between apes and other primates. While the nesting behavior of black and white ruffed lemurs, for example, appears innate [44], that of chimpanzees appears learned [45]. Accordingly, site-flexible nest construction is thought to reflect great apes’ capacity for environmental problem solving [46] and is considered by some to be the most pervasive form of material culture in great apes [5,15,17]. Thus, our results provide further evidence that an integration of the sleep quality hypothesis, sleep-site flexibility, and environmental cognition are necessary to explain early hominid evolution [41]. One remaining question is whether inter-species cognitive differences also mediate the fitness benefits of sleep quality. For example, high quality sleep may be more important for species that rely on sophisticated cognitive maps to keep track of temporal patterns of resource availability at multiple locations.

The precise nature of the tradeoffs in sleep-site selection – and the ecological conditions that favor the evolution of one strategy over another – would be further elucidated by expanding the comparisons in sleep among lemur species and between other primates. Though it was our hope that this study might help differentiate the ecological factors mediating the impact of sleep quality, the nearly negligible amount of variance in our data that can be attributed to subject species tells us there is little to be gained from attempting to explicitly model species level effects with the data at hand. This is perhaps unsurprising given the limited number of individuals per species in our study, and speaks more to a lack of statistical power than the absence of species level variation. Here, we were limited by the constraints of working with endangered species and the logistical difficulty of maintaining controlled sleep conditions in a large, multipurpose facility.

None-the-less, we can offer some insights to researchers who wish to answer these questions through the collection of additional data. We did not, for example, consider in our experimental design the important interaction of experimental condition with the order of presentation. The potential of carryover effects (the impact of one condition differentially affecting sleep during the next condition) resulting from this interaction required us to include additional parameters in our statistical model. Allowing a recovery period between experimental conditions would remove the need to statistically control for this effect, increasing the statistical power per observation without creating additional work or disturbance to the lemurs. We also take a fairly simple approach to modeling the temporal dynamics of lemur sleep due to the limited duration of our observation periods. Study of sleep patterns during extended baseline conditions would allow for more sophisticated models of baseline sleep behavior (e.g. by adding additional moving average or autoregressive parameters to account for long term homeostatic trends). These additional data would not require the time intensive and disruptive efforts of experimental manipulations, but would facilitate inference about the effects of such experiments when they are conducted.

Future work that employs these ideas may also identify other ecological co-variates of increased sleep duration in enriched sleep sites. The inclusion of smaller, fixed-point nesting species in a similar study, for example, could reveal trade-offs between more flexible sleep locations and periods of intense, high-quality sleep-in primates. Many lemurs, including those in our study, also display a high degree of cathemerality [35]; with sufficient data, it may be possible to link flexible sleep site locations with flexible sleep timing. As affordable and mobile sleep monitoring technology improves upon current methods, future research can increase the number of subjects per species used in such a study. Though we are limited in the inferences we can make about species level differences in the relationship between sleep site and sleep duration, our experimental approach has clearly demonstrated that there is a relationship in at least some lemurs. With further replication and refinement, this approach can be used to understand not just where and how primates sleep, but why they do so.

In conclusion, we found that enriched sleep-sites increase sleep duration in at least some lemurs. This highlights the importance of sleep-site conditions for lemurs, as is also known for hominoids [12,47–49] and cercopithecoids [50]. With respect to captive primate welfare, previous work has illustrated the importance of enriching sleep-sites for large brained and large bodied great apes [15,51]. In light of our findings, we suggest that managing institutions for any primate species should take care to allot resources to ensure that primates that are primed to sleep in species-specific ways, by way of environmentally modified sleep-sites, and are not using substandard sleep environments. Finally, we conclude that behaviors that influence sleep-site selection, thereby augmenting sleep quality, are evolutionarily conserved in primates and may be critically important for not only large bodied apes and monkeys, but certain prosimians as well.

## Contributorship

Vining, A.Q.: formal analysis; investigation; project administration; validation; visualization; writing – final draft preparation.

Nunn, C.L.: conceptualization; formal analysis; funding acquisition; methodology; project administration; resources; supervision; validation; writing – review & editing

Samson, D.R.: conceptualization; data curation; formal analysis; methodology; visualization; writing – original draft preparation, review & editing.

## Acknowledgments

We are grateful to the staff at the DLC and offer thanks to Erin Ehmke and David Brewer for continuous support through all aspects of this research. We thank Mark Grote for his statistical consultation and Barbara Fruth for her helpful review of this manuscript. Additionally, we would like to thank the reviewer comments that helped improve previous versions of the manuscript. This research was supported by Duke University.

## References

1. Zepelin H, Siegel JM, Tobler I. Mammalian Sleep. Princ Pract Sleep Med. 2005; 91–100. doi:10.1016/B0-72-160797-7/50015-X

2. Allison T, Cicchetti D V. Sleep in mammals: Ecological and constitutional correlates. Science (80-). 1976;194: 732–734. doi:10.1126/science.982039

3. Capellini I, Barton RA, McNamara P, Preston BT, Nunn CL. Phylogenetic analysis of the ecology and evolution of mammalian sleep. Evolution (N Y). 2008;62: 1764–1776. doi:10.1111/j.1558-5646.2008.00392.x

4. Lesku JA, Roth TC, Amlaner CJ, Lima SL. A phylogenetic analysis of sleep architecture in mammals: The integration of anatomy, physiology, and ecology. Am Nat. 2006;168: 441–453. doi:10.1086/506973

5. Fruth B, Tagg N, Stewart F. Sleep and nesting behavior in primates: A review. Am J Phys Anthropol. 2018;166: 499–509. doi:10.1002/ajpa.23373

6. Kappeler PM. Nests, tree holes, and the evolution of primate life histories. Am J Primatol. 1998;46: 7–33. doi:10.1002/(sici)1098-2345(1998)46:1<7::aid-ajp3>3.0.co;2-%23

7. Reichard U. Sleeping sites, sleeping places, and presleep behavior of gibbons (Hylobates lar). Am J Primatol. 1998;46: 35–62. doi:10.1002/(SICI)1098-2345(1998)46:1<35::AID-AJP4>3.0.CO;2-W

8. Fruth B, Hohmann G. Nest building behavior in the great apes: The great leap forward? In: McGrew WC, Marchant LF, Nishida T, editors. Great Ape Societies. Cambridge University Press; 1996. pp. 225–240.

9. McGrew WC. The Cultured Chimpanzee. Cambridge: Cambridge University Press; 2004. doi:10.1017/cbo9780511617355

10. Samson DR. The chimpanzee nest quantified: Morphology and ecology of arboreal sleeping platforms within the dry habitat site of Toro-Semliki Wildlife Reserve, Uganda. Primates. 2012;53: 357–364. doi:10.1007/s10329-012-0310-x

11. Videan EN. Sleep and sleep-related behaviors in Chimapnzee (Pan troglodytes). Miami University. 2005.

12. Stewart FA, Pruetz JD, Hansell MH. Do Chimpanzees Build Comfortable Nests. Am J Primatol. 2007;69: 930–939. doi:10.1002/ajp

13. Cheyne S, Rowland D, Höing A, Husson S. How orang-utans choose where to sleep: comparison of nest site variables. Asian Primates J. 2013;3: 13–17.

14. Samson DR, Shumaker RW. Species differences in sleep quality between captive orangutans (Pongo pygmaeus) and baboons (Papio papio). American journal of Physical Anthropology. 2014. p. 216. doi:10.1002/ajpa/22488

15. Shumaker RW, Samson DR. Documenting orang-utan sleep architecture: Sleeping platform complexity increases sleep quality in captive Pongo. Behaviour. 2013;150: 845–861. doi:10.1163/1568539X-00003082

16. Anderson JR. Sleep-related behavioural adaptations in free-ranging anthropoid primates. Sleep Med Rev. 2000;4: 355–373. doi:10.1053/smrv.2000.0105

17. McGrew WC. Chimpanzee Material Culture: Implications for Human Evolution. Cambridge University Press; 1992.

18. Samson DR, Vining A, Nunn CL. Sleep influences cognitive performance in lemurs. Anim Cogn. 2019;22: 697–706. doi:10.1007/s10071-019-01266-1

19. Huebner F, Fichtel C, Kappeler PM. Linking cognition with fitness in a wild primate: Fitness correlates of problem-solving performance and spatial learning ability. Philos Trans R Soc B Biol Sci. 2018;373. doi:10.1098/rstb.2017.0295

20. Bray J, Samson DR, Nunn CL. Activity patterns in seven captive lemur species: Evidence of cathemerality in Varecia and Lemur catta? Am J Primatol. 2017;79: 1–9. doi:10.1002/ajp.22648

21. Stone KL, Ancoli-Israel A. Actigraphy. Fifth. In: Kryger MH, Roth T, William DC, editors. Principles and Practices of Sleep Medicine. Fifth. St. Louis, Missouri: Saunders; 2010. pp. 1668–1675.

22. Andersen ML, Diaz MP, Murnane KS, Howell LL. Effects of methamphetamine self-administration on actigraphy-based sleep parameters in rhesus monkeys. Psychopharmacology (Berl). 2013;227: 101–107. doi:10.1007/s00213-012-2943-2

23. Howell L. Reply to Sleep parameters in rhesus monkeys by using Actigraphy. Psychopharmacology (Berl). 2013;228: 511. doi:10.1007/s00213-013-3171-0.Reply

24. Barrett CE, Noble P, Hanson E, Pine DS, Winslow JT, Nelson EE. Early adverse rearing experiences alter sleep-wake patterns and plasma cortisol levels in juvenile rhesus monkeys. Psychoneuroendocrinology. 2009;34: 1029–1040. doi:10.1016/j.psyneuen.2009.02.002

25. Sri Kantha S, Suzuki J. Sleep quantitation in common marmoset, cotton top tamarin and squirrel monkey by non-invasive actigraphy. Comp Biochem Physiol - A Mol Integr Physiol. 2006;144: 203–210. doi:10.1016/j.cbpa.2006.02.043

26. Zhdanova I V., Geiger DA, Schwagerl AL, Leclair OU, Killiany R, Taylor JA, et al. Melatonin promotes sleep in three species of diurnal nonhuman primates. Physiol Behav. 2002;75: 523–529. doi:10.1016/S0031-9384(02)00654-6

27. Samson DR, Bray J, Nunn CL. The cost of deep sleep: Environmental influences on sleep regulation are greater for diurnal lemurs. Am J Phys Anthropol. 2018;166: 578–589. doi:10.1002/ajpa.23455

28. Campbell SS, Tobler I. Animal sleep: A review of sleep duration across phylogeny. Neurosci Biobehav Rev. 1984;8: 269–300. doi:10.1016/0149-7634(84)90054-X

29. Kawada T. Sleep parameters in rhesus monkeys by using actigraphy. Psychopharmacology (Berl). 2013;228: 509. doi:10.1007/s00213-013-3170-1

30. Pinheiro J, Bates D. Extending the Basic Linear Mixed-Effects Model. Mixed-Effects Models in S and S-PLUS. Springer Science & Business Media; 2006. doi:10.1007/b98882

31. Pinheiro J, Bates D, DebRoy S, Sarkar D. R Development Core Team. 2010. nlme: Linear and Nonlinear Mixed Effects Models. R Packag version 3. 2007; 1–97.

32. Team RC. R: A Language and environment for statistical computing. Vienna, Austria: R Foundation for Statistical Computing; 2020.

33. Anderson JR. Sleep, sleeping sites, and sleep-related activities: Awakening to their significance. Am J Primatol. 1998;46: 63–75. doi:10.1002/(SICI)1098-2345(1998)46:1<63::AID-AJP5>3.0.CO;2-T

34. Karpanty SM, Wright PC. Predation on Lemurs in the Rainforest of Madagascar by Multiple Predator Species: Observations and Experiments. Primate Anti-Predator Strateg. 2007; 77–99. doi:10.1007/978-0-387-34810-0_4

35. Colquhoun IC. Predation and cathemerality: Comparing the impact of predators on the activity patterns of lemurids and ceboids. Folia Primatol. 2006;77: 143–165. doi:10.1159/000089701

36. Kappeler PM, Nunn CL, Vining AQ, Goodman SM. Evolutionary dynamics of sexual size dimorphism in non-volant mammals following their independent colonization of Madagascar. Sci Rep. 2019;9. doi:10.1038/s41598-018-36246-x

37. Day RT, Elwood RW. Sleeping site selection by the golden-handed tamarin Saguinus midas midas: The role of predation risk, proximity to feeding sites, and territorial defence. Ethology. 1999;105: 1035–1051. doi:10.1046/j.1439-0310.1999.10512492.x

38. Baden AL. A description of nesting behaviors, including factors impacting nest site selection, in black-and-white ruffed lemurs (Varecia variegata). Ecol Evol. 2019;9: 1010–1028. doi:10.1002/ece3.4735

39. Nunn CL, Altizer S. Infectious Diseases in Primates. New York: Oxford University Press; 2006.

40. Nunn CL, Heymann EW. Malaria infection and host behavior: A comparative study of Neotropical primates. Behav Ecol Sociobiol. 2005;59: 30–37. doi:10.1007/s00265-005-0005-z

41. Samson DR, Nunn CL. Sleep intensity and the evolution of human cognition. Evol Anthropol. 2015;24: 225–237. doi:10.1002/evan.21464

42. Nunn CL, Samson DR. Sleep in a comparative context: Investigating how human sleep differs from sleep in other primates. Am J Phys Anthropol. 2018;166: 601–612. doi:10.1002/ajpa.23427

43. Samson DR, Crittenden AN, Mabulla IA, Mabulla AZP, Nunn CL. Chronotype variation drives night-time sentinel-like behaviour in hunter – gatherers. Proc R Soc B Biol Sci. 2017;284. doi:10.1098/rspb.2017.0967

44. Pereira ME, Klepper A, Simons EL. Tactics of care for young infants by forest-living ruffed lemurs (Varecia variegata variegata): Ground nests, parking, and biparental guarding. Am J Primatol. 1987;13: 129–144. doi:10.1002/ajp.1350130204

45. Videan EN. Bed-Building in Captive Chimpanzees (Pan troglodytes): The Importance of Early Rearing. Am J Primatol. 2006;68: 745–751. doi:10.1002/ajp

46. McGrew WC. Tool-use by free-ranging chimpanzees: the extent of diversity. J Zool. 1992;228: 689–694. doi:10.1111/j.1469-7998.1992.tb04469.x

47. Koops K, McGrew WC, de Vries H, Matsuzawa T. Nest-Building by Chimpanzees (Pan troglodytes verus) at Seringbara, Nimba Mountains: Antipredation, Thermoregulation, and Antivector Hypotheses. Int J Primatol. 2012;33: 356–380. doi:10.1007/s10764-012-9585-4

48. Samson DR, Hunt KD. A Thermodynamic Comparison of Arboreal and Terrestrial Sleeping Sites for Dry-Habitat Chimpanzees (Pan troglodytes schweinfurthii) at the Toro-Semliki Wildlife Reserve, Uganda. Am J Primatol. 2012;74: 811–818. doi:10.1002/ajp.22031

49. Stewart FA. Brief communication: Why sleep in a nest? empirical testing of the function of simple shelters made by wild chimpanzees. Am J Phys Anthropol. 2011;146: 313–318. doi:10.1002/ajpa.21580

50. Bert J, Balzamo E, Chase M, Pegram V. The sleep of the baboon, Papio papio, under natural conditions and in the laboratory. Electroencephalogr Clin Neurophysiol. 1975;39: 657–662. doi:10.1016/0013-4694(75)90079-6

51. Shumaker R. Gorilla Enrichment. In: Ogden J, Wharton D, editors. Management of Gorillas in Captivity. Atlanta: Gorilla species survival plan and the Atlanta/Fulton County Zoo; 1997. pp. 102–110.

